# Species climate-niche properties suggest that both physiological tolerance and stress dominance shape plant community assembly across a tropical dry ecosystem

**DOI:** 10.64898/2026.07.28.741168

**Authors:** Swastik Patnaik, Souparna Chakrabarty, Reshma M. Ramachandran, Parth Sarathi Roy, Meghna Krishnadas

## Abstract

Climate influences community assembly by constraining the conditions under which species can persist, but the consistency of macroecological processes across different ecosystems remains unclear. Interspecific variation in climatic niche properties that govern community assembly across climate gradients can be gleaned from species distributions. Species-climate associations, niche properties, and resulting assembly remains poorly understood for plants in tropical dry ecosystems, and among different life-forms.

We examined how species-climate associations governed community assembly of four plant life-forms across 60,000 km^2^ of tropical dry ecosystem in peninsular India. In the 1750 km long Eastern Ghats mountain range, we surveyed vegetation in 2500 20 x 50 m plots. Across the north-south climatic gradient from cool, wet and seasonal sites to warm, dry and less-seasonal sites, from presence-absence data of 1601 species in four life forms (trees, shrubs, herbs, climbers), we performed Bayesian Hierarchical Modelling of Species Communities to predict conditions of occurrence for each species. From this, we derived species’ climatic niche optima and niche width, and examined the local-level prevalence of niche properties across the climate gradient to glean assembly.

Most tree, shrub and climber species associated with warm-dry conditions had wider niches than species associated with cool-wet conditions, which also resulted in assemblages having wider niches at warmer sites. Herbaceous species, by contrast, had narrower niches when associated with warmer conditions, where narrow-niche species dominated the assemblage. In all life-forms, however, assemblages in both warm-dry and cool-humid conditions consisted of species primarily associated with and preferring (having their optima in) those conditions, suggestive of stress dominance. Further evidence for stress dominance in shrubs and climbers came from assemblages in the warmest and coolest sites having low richness and comprising mainly of species with wider niches. Tree richness increased at either end of the gradient, while herb richness increased in warm-dry conditions.

Overall, our results suggest that plant life-forms differ in the processes driving assembly across broad climate gradients in a tropical dry ecosystem, but many herbs were specialized to warm-dry climates. To understand life-form-specific responses to environmental gradients in tropical dry ecosystems, future work should incorporate traits and evolutionary history.

## 1. INTRODUCTION

Understanding why species occur where they do is a fundamental question in biogeography and macroecology. Species distributions in relation to abiotic and biotic factors offer insights into drivers and mechanisms of community assembly at multiple scales, and such efforts have gained renewed interest with the urgent need to predict species responses to climate change (Murphy & Smith, 2021). Aided by increased availability of environmental and biodiversity data and recent advances in modelling techniques (Tikhonov et al., 2020), a better understanding of how climate shapes and constrains communities can reveal macroecological drivers of biodiversity patterns (Walter et al., 2020), understand variation in biodiversity-ecosystem function relationships, and identify biodiversity reserves (Noroozi et al., 2019) to inform conservation planning (Dullinger et al., 2017).

Species distributions in relation to climate reveal properties of their broad-scale niches (Takola & Schielzeth, 2022). While niche width represents the range of realized conditions where a species occurs, species usually achieve peak performance and maximize fitness in a narrower set of conditions, thought of as species’ niche optima (Lian, et al., 2022). These niche properties (width and optima) shape community assembly across climate gradients. For example, in multiple tropical landscapes, tree species’ association with precipitation regulated community composition across large spatial scales (Condit et al., 2013; Esquivel-Muelbert et al., 2017; Krishnadas et al., 2021). Wet-associated species are often restricted in their distribution by the physiological challenges of dry conditions, and thereby tend to have narrower niches than dry-associated species (Baltzer et al., 2008; Baltzer & Davies, 2012), which can persist in dry and wet sites. This leads to communities being composed of dry-associated species in areas with more pronounced seasonal drought whereas both wet- and dry-associated species co-occur in less seasonal conditions resulting in higher diversity than at drier sites (Krishnadas et al., 2021). Two major hypotheses may explain distribution patterns and climate niches: the physiological tolerance hypothesis (Larcher, 2003) and stress-dominance hypothesis (Díaz et al., 1998). The physiological tolerance hypothesis predicts that abiotic filtering based on species’ tolerance limits will generate nested distributions from benign to harsh environments (e.g., Esquivel-Muelbert et al., 2017). This would be the basis for dry-tolerant species persisting in wet conditions but not *vice versa*. In contrast, the stress-dominance hypothesis proposes that abiotic stress primarily structures communities under harsh conditions, whereas biotic interactions such as competition impose limitations under more benign conditions (Kuczynski & Grenouillet, 2018). Stress-dominance may lead to disjunct distributions such that species adapted to stressful climates get outcompeted from benign climates due to potential trade-offs with competitive ability.

Furthermore, species-climate linkages among plants can vary among plant life forms, but comparative studies are few (Yang et al., 2025), limiting our understanding of whether different life forms respond similarly to climatic gradients and environmental change. Trees have larger and deeper root systems than shrubs or herbs to access water from deeper soil layers (Kuhnhammer et al., 2023). Some tree species can also store water in their stems and roots (Borchert, 1994) which may allow them to persist across a wide range of climatic conditions. Climbers can also have deep roots or draw their water from host trees (Ghazoul & Sheil, 2010), helping them occupy large climatic gradients. Shrubs, by contrast, are typically shallow rooted and do not have the capacity for stem water storage (Becker & Castillo, 1990), especially those adapted to shade, and may be constrained to specific microenvironments in dry ecosystems (Mills & Schwartz, 2025). Herbaceous species may prefer warmer, drier sites and can be limited by shaded understory in wetter sites as canopies become thicker than drier sites (Rheault et al., 2015; Moser et al., 2004).

Despite the large and growing body of work on the relationships between climate and species distributions, some biomes remain understudied, such as Tropical Dry Ecosystems (Norden et al., 2026; henceforth TDE). TDE are defined by extended periods of severe water deficit conditions, accompanied by warm temperatures and high evapotranspiration rates (Balvanera, Quijas, & Pérez-Jiménez, 2011; Davies et al., 2023). While characterized by strong seasonality of precipitation, TDEs exist across gradients of temperature and precipitation conditions to which species may respond differentially (Méndez-Toribio et al., 2016). The outstanding question is whether TDE species having been filtered based on their suitability to regional conditions are generalists that persist across most sites, or vary in their niche properties within the broader climate space of TDE.

Evidence suggests that plant species in TDE may be operating at their thermal limits. Hence, even relatively short gradients in temperature may regulate species occurrences across space (Norden et al., 2026). Similarly, while TDE species may generally withstand seasonal drought, differential moisture requirements may regulate species distributions across prevailing moisture gradients (Rajendra et al. in revision). Understanding interspecific variation to climatic factors and how species-climate associations shape compositional patterns is an essential first step in predicting changes to tree communities in TDE with future warming and drought (Rodrigues et al., 2015).

We sought to understand species-climate associations and its role in structuring plant communities across broad macroclimate gradients in the tropical dry ecosystem of Eastern Ghats, a 1750 km long mountain chain along the east coast of peninsular India (Panda et al., 2020). In the Indian subcontinent, there have been landscape-level studies of species-climate associations in the Western Ghats (Gopal et al., 2023; Krishnadas et al., 2021) and the Himalayas (Manish et al., 2016). In contrast to these wet forests, the Eastern Ghats is a typical TDE with drier and more seasonal climate where the vegetation comprises tropical moist and dry deciduous forests through to savanna (Gandhi & Sundarapandian, 2020) (Anupama et al., 2000) that harbour many endemic plant species (Ramachandran et al., 2020). Assessments of species distributions in the Eastern Ghats have either focused on selected species (Remya et al., 2015) or done at small spatial scales (Behera et al., 2020), which do not reveal how climatic conditions drive community assembly across the regional gradient of climate. To understand vegetation assembly across a ∼60000 sq.km of TDE in peninsular India, we asked:

1. How does climate shape species distributions and do patterns of species-climate associations vary by life form (trees, shrubs, climbers, herbs)?

We expected some species to strongly associate with cooler-wetter sites or with warmer-drier sites (specialists) while many were expected to be generalists that occur widely across the climate gradient. Species associated with warmer-drier sites were expected to persist in cooler-wetter sites, but not vice-versa. Among life forms, we expected trees and climbers to have more generalists while shrubs and herbs prefer wetter and drier sites respectively.

2. Does climate niche width and optimal niche conditions vary among species and life forms?

We expected that species associated to or having their optima at the extremes of cool-wet and hot-dry regions might be more specialized and have a smaller niche width than species associated to or peaking at intermediate conditions.

3. Do species-climate associations suggest community assembly via physiological tolerance or stress dominance and do signals of mechanisms differ by life form?

We hypothesized that the hotter drier regions are likely to be dominated by dry-associated species, but cool-wet regions can support species associated with hot-dry and cool-wet conditions.

## 2. MATERIALS AND METHODS

### 2.1 Study site

The Eastern Ghats mountain range stretches across 1,750 km along the eastern coast of peninsular India encompassing areas within the states of Odisha, Telangana, Andhra Pradesh, Karnataka and Tamil Nadu. The geographic extent of the Eastern Ghats is defined from the Similipal hills to the north of the Mahanadi river in the north to the Vaigai river in the south (Prasad et al. 2019). The study site encompasses various ecoregions such as Savanna woodland, East Deccan dry evergreen forests, South Deccan Plateau dry deciduous forests, and Eastern Highlands moist deciduous forests. Climatically, the region is characterized by strong seasonality, with pronounced dry periods, high temperatures, and variable precipitation, making it representative of TDE across the globe. These ecosystems experience high evapotranspiration and periodic water deficit, which are critical drivers of vegetation distribution and composition.

### 2.2 Data collection

The dataset originates from surveys conducted for the Manual of Biodiversity Characterisation at Landscape Level using Remote Sensing & Geographical Information System, published by the Department of Space and the Department of Biotechnology (Roy et al., 2013). A digital map of the vegetation types was obtained through GIS tools. Based on this information, ground surveys were conducted to establish 2561 GPS tagged field plant inventory plots of 20 x 50 m size, using stratified modified Whittaker plot method (Stohlgren et al. 1995) where plots were placed to encompass maximum landscape heterogeneity. At each plot all the species of whose adult plants were present were recorded. In total, 1601 species have been recorded, with 639 herbs, 311 shrubs, 209 climbers and 442 tree species.

### 2.3 Environmental variables and climatic gradients

We obtained the environmental variables for each plot from the WorldClim version 2.1 (Fick & Hijmans, 2017) database which has global climatic data at ∼1 km (2.5 arcsec) spatial resolution. We used the standard 19 BioClim variables which incorporate temperature, precipitation and seasonality metrics. We also included Climatic Water Deficit (CWD) as an ecologically meaningful indicator of water stress, obtained from TerraClimate-<u>Climatology Lab</u>, (Abatzoglou et al., 2018). Prior to analysis, environmental variables were transformed using a custom pre-processing routine to ensure comparability across predictors. To reduce dimensionality and account for multicollinearity among climatic variables, we performed a principal component analysis (PCA) using the ‘prcomp’ function in R with variables centred and scaled. The PCA was conducted on this set of 20 climatic variables describing annual, seasonal, and extreme temperature and precipitation conditions. The first three principal components (PC1–PC3) were used as predictors in the JSDM models and subsequent downstream analyses. PC1 primarily represented a gradient from cool–wet to warm–dry conditions (Figure 1) while PC2 and PC3 captured gradients related to seasonality and extreme climatic variability.

**FIGURE 1.**
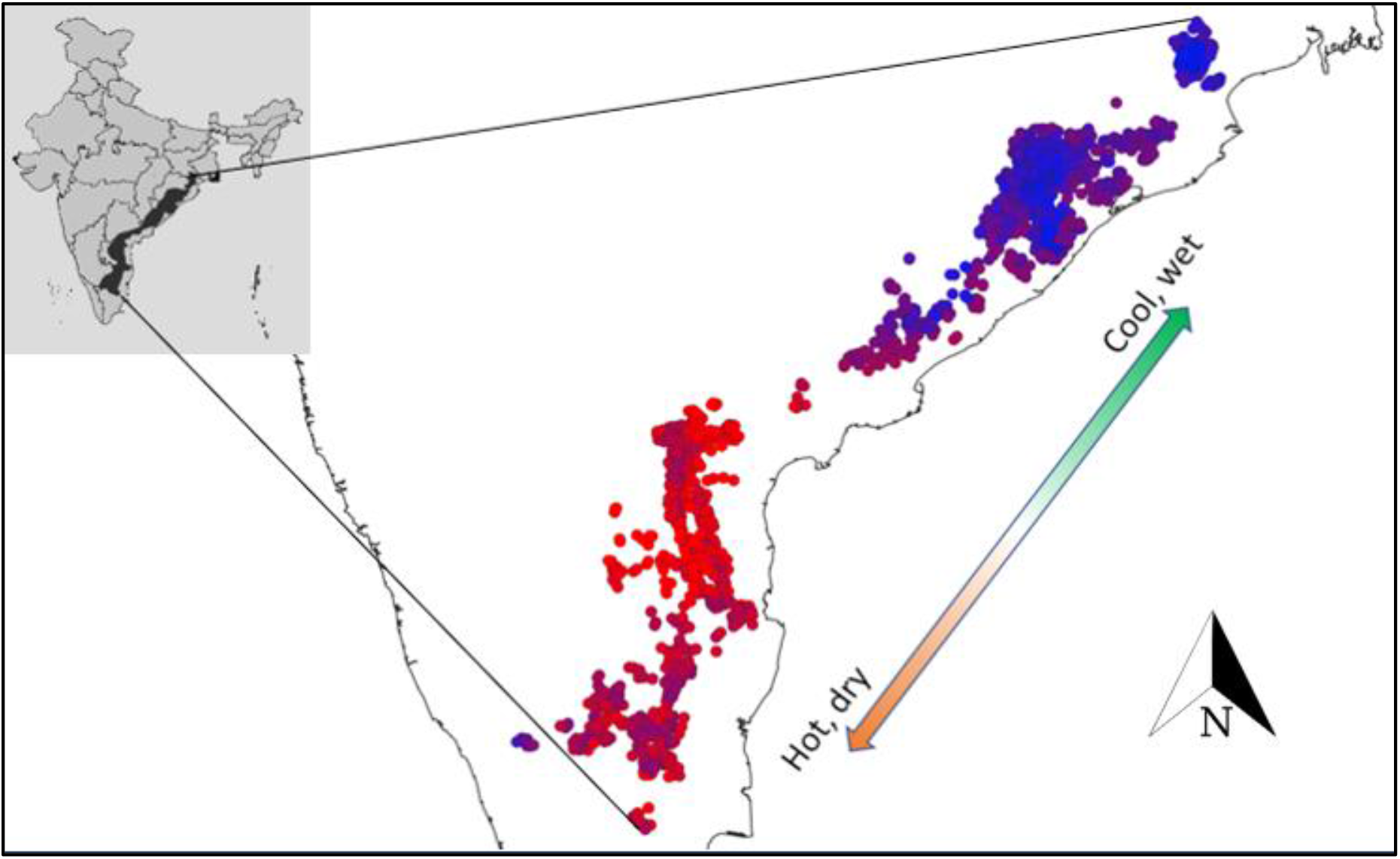
Locations sampled across the Eastern Ghats. Vegetation was sampled in 2500 plots all along the Eastern Ghats mountain range in peninsular India, covering the full spectrum of climatic gradient from cool, wet regions in the North to hot and dry regions of the south. Each point represents a 10×10 m plot where species identity was recorded. The shading of points represents the climatic gradient along the principal component (PC1) axis, with blue shades representing cooler, wetter climate and red shades representing warmer, drier climate.

### 2.4 Statistical analysis

#### Hierarchical modelling of species communities

The presence-absence data for each species in each plot was transformed into a binary matrix with *A_ij_* = 1 representing species *i* being present in plot *j* and *A_ij_* = 0 similarly representing absence. Species occurrences were modelled using the Hierarchical Model of Species Communities (HMSC) framework (Ovaskainen et. al. 2017). HMSC is a Bayesian joint species distribution modelling framework that allows simultaneous estimation of species responses to different drivers, accounting for co-variation among species. We fit separate models for climate and topography, where species occurrences were modelled as a function of the first three climatic principal components (PC1–PC3) or as a function of slope and aspect of each plot, respectively for each life form. We modelled species distributions using a generalized linear mixed model (GLMM) framework as implemented in HMSC, incorporating fixed environmental effects and random effects to account for spatial and hierarchical structure, with species responses assumed to follow a binomial error distribution linked to predictors via a logit link function. Models included quadratic terms to account for nonlinear responses of species, with species-specific intercepts and slopes estimated per predictor. Models were formulated as:

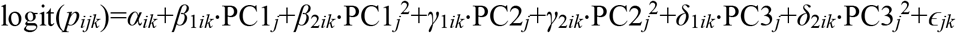

for climatic models and,

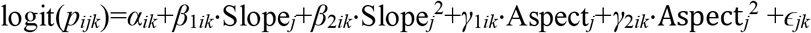

for topographic models where, *pijk* is the probability of occurrence for species *i* of life form *k* at site *j*, *αik* is the species-specific intercept within life form *k*, *β, γ, δ* are species-specific coefficients for each predictor, and *ɛjk* are plot-level random effects accounting for spatial autocorrelation.

Only species that occurred in more than 15 plots were considered for modelling, which resulted in 193 tree, 92 shrub, 136 herb and 64 climber species in the analysis. Models were fitted using Gibbs Markov Chain Monte Carlo (MCMC) sampling. Each model was sampled to obtain 1000 posterior samples, retaining every 100th iteration. Convergence and model performance were evaluated using standard MCMC diagnostics. Convergence of species-specific regression coefficients was assessed using the potential scale reduction factor (PSRF), with values ≤1.1 considered indicative of adequate convergence. Diagnostic routines were applied separately to each life-form and environmental model.

We used the fitted models to extract posterior estimates of species–environment associations. Positive and negative values of the first-order slopes indicated whether species increased or decreased from low to high values of the climate gradient. The second order coefficient indicates the curvature of the response, allowing us to infer whether species exhibited nonlinear responses with an optimum. Using these estimates, we predicted probabilities of species occurrence for each plot. Predicted occurrences were converted to binary presence–absence predictions using a probability threshold of 0.01, and these predictions were used to estimate species niche properties across the climate gradient.

### 2.5 Quantifying species’ environmental niche

We used predicted occurrence probabilities for each species across each climate gradient to identify locations occupied with a non-zero probability based on the 95% credible intervals of estimates using the first two principal components of our PCA as environmental variables which together explain 79% of the environmental variation across the system. We estimated species niche optima as the centroid of occupied points in PC1-PC2 space derived from a Voronoi tessellation (Du et al., 1999). Niche width was calculated as the area of the species’ minimum convex polygon (convex hull) in PC1-PC2 space, standardised by the total environmental space available in the dataset.

We first analysed patterns of species’ climate associations and how they vary by life form. We look at relationships of climate association, i.e., posterior first-order coefficient, with niche optima and niche width to find how patterns of association change across species present in locations with different climatic conditions for each life form. For our second question, relationships between species’ climatic associations (first-order coefficient), niche optima, and niche widths were used to infer whether species with different climatic conditions show differing niche properties and whether this differs across life forms. These relationships were analysed using generalized additive models (GAMs) with life form as a categorical variable.

### 2.6 Community-level responses along climatic gradients

To address our third question regarding how species’ niche properties shape community assembly, species-level climate association and niche properties were aggregated to the plot level to examine community-level prevalence of niche properties. For all species occurring in each plot, we calculated mean niche width and mean species’ climate association (mean of species-level first-order regression coefficient), estimated from their predicted distribution. Species were considered present at a site if the 95% CrI of their probability of occurrence did not overlap zero. To assess whether community-level patterns were sensitive to the way plots were aggregated along the climatic gradient, plots were grouped into bins based on their position along the climate axis, and species-level metrics were averaged across all species occurring in all plots within each bin. This allowed us to estimate how community-level niche properties and climate associations vary systematically along the environmental gradient, while reducing noise from individual plot-level variation. Results remained similar upon repeating the site-level assessments using species’ presence at a site regardless of the credible intervals overlapping zero, so we retained the more statistically robust and conservative estimates based on 95% CrI.

Community-level properties were analysed along the primary climatic gradient (PC1) using generalized additive models (GAMs), with PC1 as a predictor and life form included as a categorical variable interacting with PC1 to allow life form–specific responses. Plot identity was included as a random intercept. Analyses were restricted to PC1 because it explained 54.3% of the total climatic variation, more than twice that explained by the second principal component. To visualize trends along the climatic gradient, plots were grouped into quantile bins based on their PC1 values. Mean community-level responses within each bin were then calculated and smoothed using GAMs. Finally, predicted species occurrences were used to estimate expected patterns of species richness and niche properties across the climatic gradient. Specifically, we tested whether hotter and drier communities exhibited lower species richness and narrower community-level niche widths. Lower richness at limits of the climatic gradient would be consistent with stronger environmental filtering, where species adapted to harsh conditions can also occur under benign conditions, but species adapted to benign conditions are excluded from harsher environments. In contrast, lower niche widths at climatic range limits would indicate communities dominated by climatically restricted species, consistent with increasing specialization.

## 3. RESULTS

### 3.1 Species distributions across climate gradients

Mean temperature of the driest quarter followed by precipitation in the warmest quarter mainly differentiated sites across the Eastern Ghats TDE (Supplementary Information, Figure S1 (a)), characterized by a shift from cool, humid to warmer, drier but less seasonal conditions (more consistently dry and warm) as we go southwards. Overall, 85% of species (414 out of 485) showed significant responses to the climatic gradient. Among these, 58% (277 species) were associated with conditions of hotter drier regions (warm associated) and 27% (137 species) percent with cooler wetter regions (cool associated), while only 15% (71 species) did not show any climatic association, i.e., their distribution was agnostic to climatic variation across the study area, indicating ecological generalists. These patterns did not differ significantly among life forms (F = 1.786, p = 0.1515), even as the proportion of warm associated species varied from 62% for climbers to 52% for trees (Supplementary Information S1, Figure S4). The consistent patterns across different life forms underscores the role of climate as a filter on assembly of vegetation communities.

### 3.2 Species’ climate-niche properties

Niche optima, defined as the centroid of occupied points within PC1-PC2 space, varied among species along the climate gradient captured by PC1. Species associated with warmer climates (positive values of PC1) had niche optima in warm-dry regions, whereas those associated with cooler climates attained their optimum in cool-wet regions (Figure 2a, Table 1). While this pattern was generally consistent for life forms, trees demonstrated the most pronounced shift in niche optima, with a rapid transition from cool- to warm-associated species followed by a plateau under warmer conditions (Table 1). In other words, a majority of warm-associated tree species had similar niche optima, suggesting similar distribution patterns within the available climate space.

**FIGURE 2.**
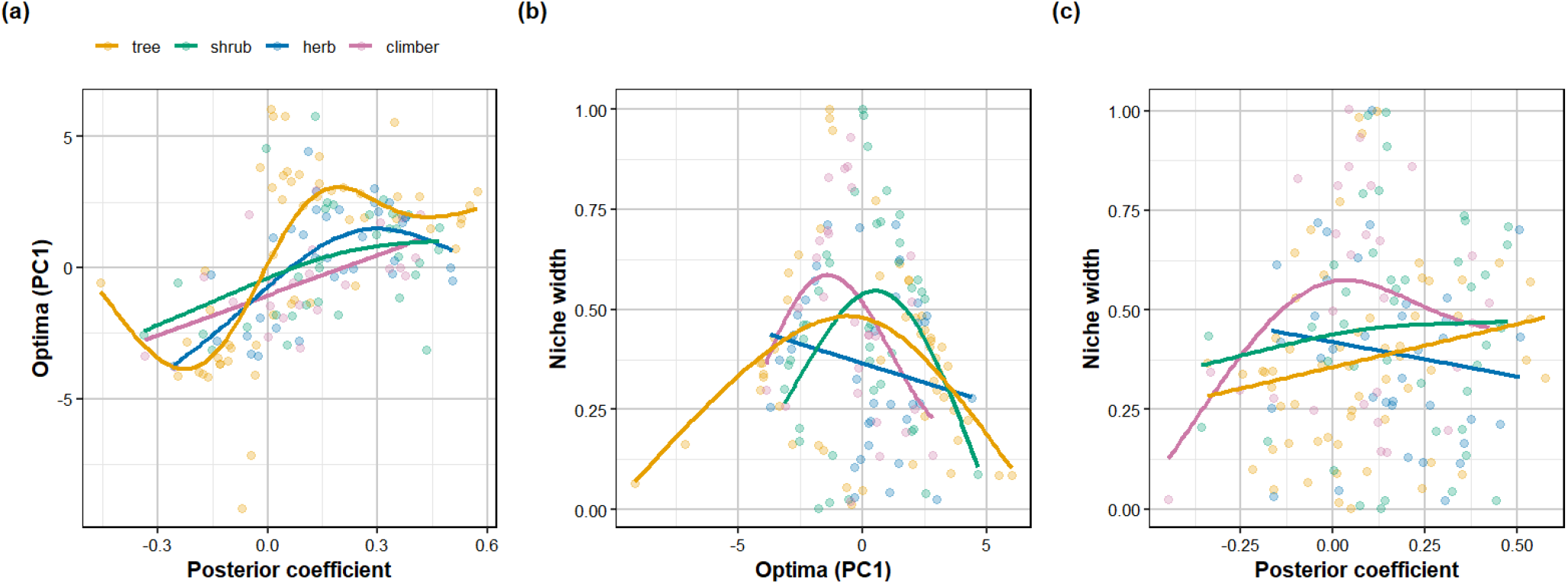
Patterns of species’ niche properties. Relationships between (a) species niche optima (weighted centroid of occupied climate values) and (posterior) coefficient of association to PC1, i.e., directional response to climate in the probability of occurrence (b) niche width and niche optima and (c) niche width and posterior coefficient. In all life forms, a large proportion of species had associations towards warm-dry conditions (posterior coefficient > 0).

**TABLE 1.**
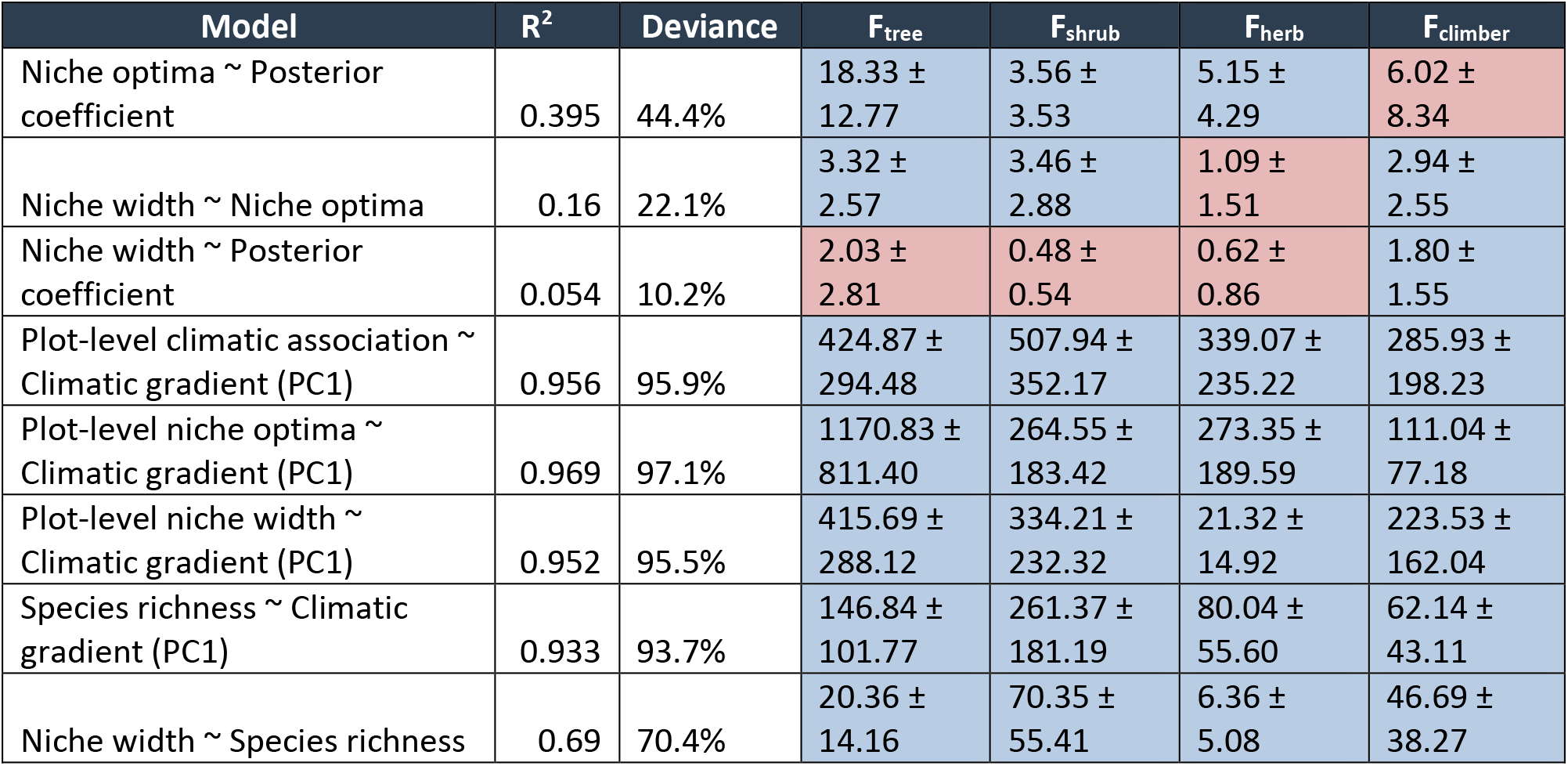
Summary of GAM statistics for all models. Values are presented as R², deviance explained (%), and F-statistics for each lifeform (tree, shrub, herb, climber) along with 95% confidence intervals. Higher F-values indicate stronger nonlinear relationships. Confidence intervals were approximated using the standard error derived from the effective degrees of freedom of the smooth terms (SE = F / √(2 × Ref.df)), with 95% CIs calculated as F ± 1.96 × SE. Significant relationships are highlighted in blue and non-significant in red.

Signals of specialization were revealed in the nonlinear relationship of species niche width against optima (Figure 2b, Table 1). Species with peak occurrence at either cool-wet or warm-dry peripheries of the climate gradient tended to have narrower niches compared to species preferring intermediate climatic conditions exhibited broader niches, suggesting greater ecological flexibility. Herbs did not show a statistically significant pattern. However, patterns of niche width against posterior climate associations (Figure 2c, Table 1) depict a general pattern, albeit weak, towards increasing at high values of the posterior coefficient. Overall, these findings indicate that ecological specialization increases toward climatic limits, highlighting the strength of environmental filtering under both cool-wet and warm–dry conditions.

### 3.3 Community assembly across the climatic gradient

Along the climatic gradient from cool-wet to warm-dry sites, mean plot-level species associations showed that assemblages clustered into species associated with and with their optima towards either of the climate conditions, for all life forms (Figure 3a, 3b; Table 1). Tree, shrub, and climber assemblages at the cool-wet parts of the climate gradient comprised of narrow-niche species whereas assemblages at the warm-dry parts comprised of species with wider niches (Figure 3c; Table 1). However, mean niche width of tree assemblages declined towards the warmest sites. By contrast, herb assemblages at cool-wet sites had species with wider niches whereas warm-dry sites comprised of narrow niche species (Figure 3c, Table 1).

**FIGURE 3.**
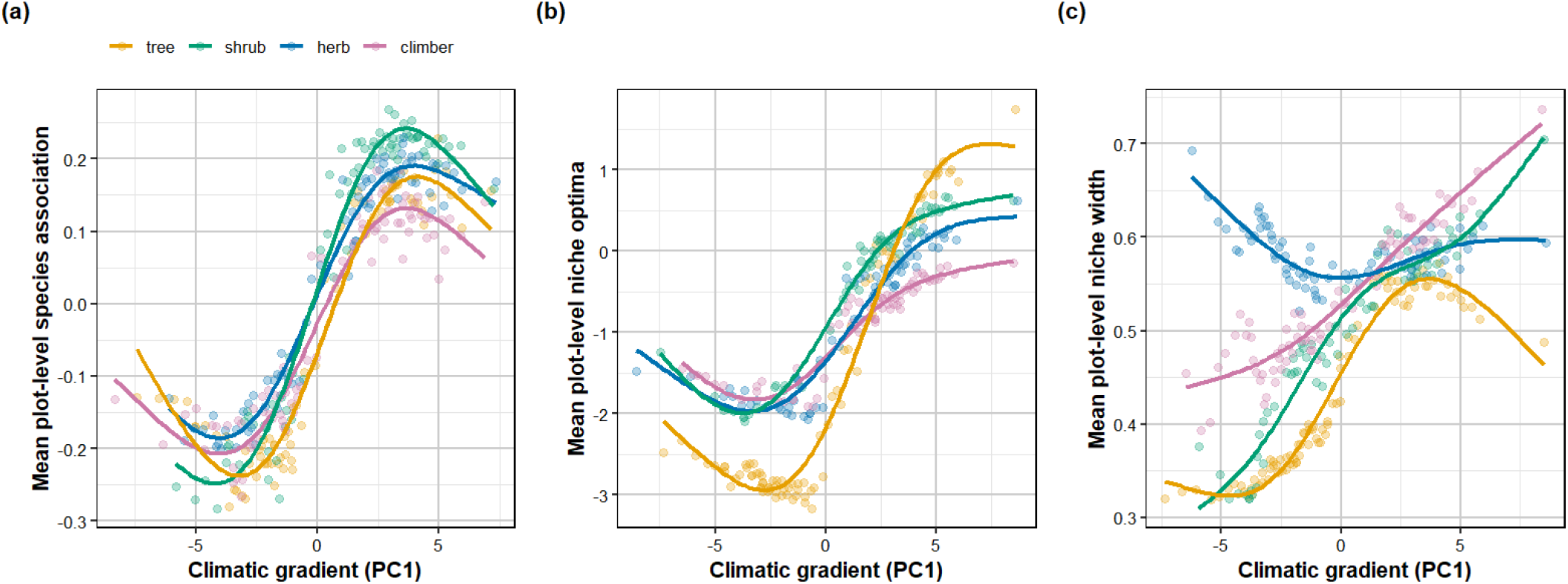
Plot-level niche properties across the climatic gradient. For all species predicted to occur at a climate value, (a) mean of the first-order regression coefficient (climate association), (b) mean niche optima, and (c) mean niche width. Patterns were assessed in relation to the dominant climatic gradient (PC1). Species’ occurrences at a climate value were predicted from Bayesian joint species’ distribution models.

### 3.4 Richness patterns

For estimated species richness across the climate gradient, trees showed a different trend from other life forms (Table 1). Predicted tree richness was lower at the intermediate climate conditions compared to warmer and cooler ends. By contrast, predicted richness of shrubs, climbers and herbs peaked towards warmer, drier sites but then declined at the warmest sites (Figure 4a, Table 1). Furthermore, in shrub and climber communities, species-rich plots had higher average plot-level niche width, suggesting that species richness may increase with the presence of generalist species. Herbs showed the opposite trend: species-rich assemblages largely comprised of species with narrow niches, indicating that each end of the climate gradient seems to have wide ranging communities, especially the wetter sites, which suggests that herbs in drier sites are likely to be specific to those conditions. Tree communities showed no clear trends (Figure 4b, Table 1).

**FIGURE 4.**
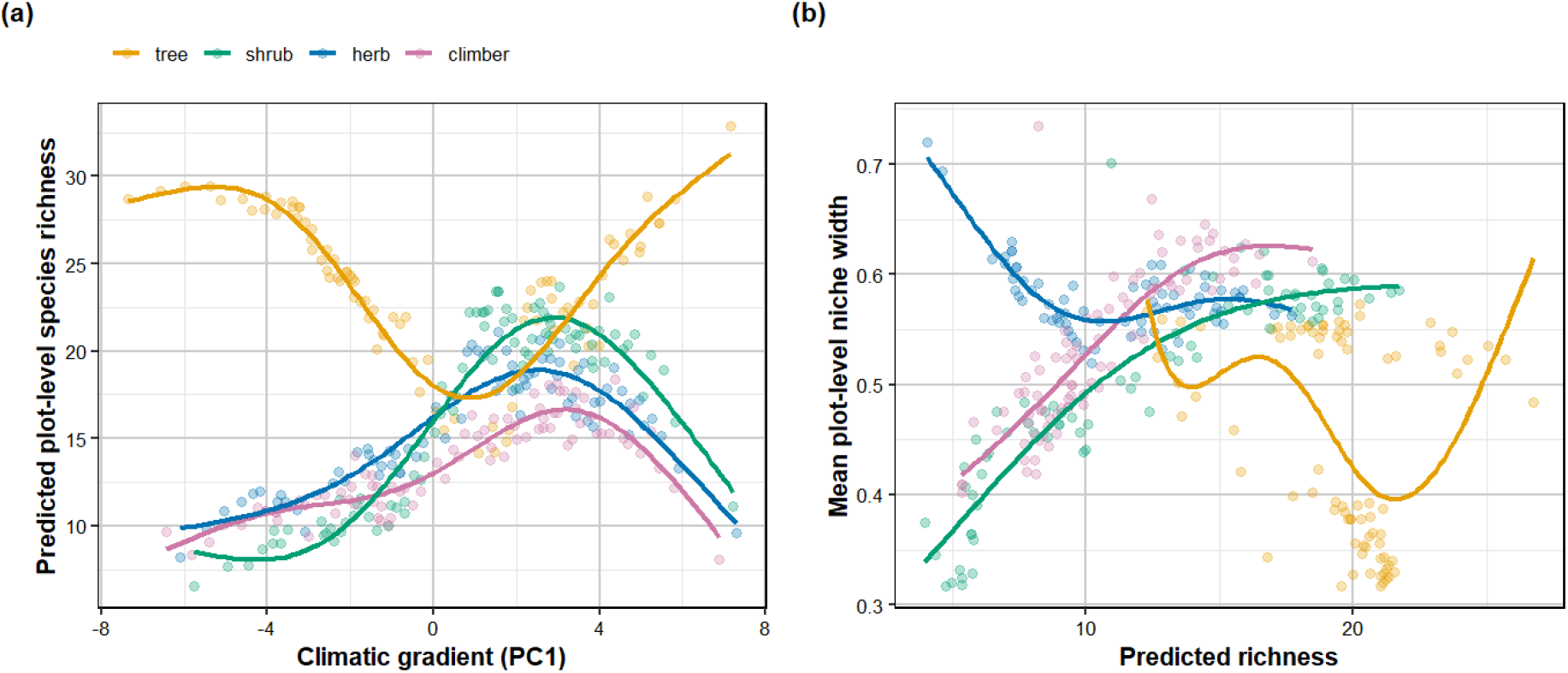
Community-level species richness across the climatic gradient. (a) Mean plot-level species richness estimated as the average number of species at each percentile of the climatic gradient, based on predicted occurrences from the posteriors of Bayesian joint species distribution models plotted against the values of PC1 (climatic gradient) for each plot and (b) mean plot-level niche width of the community against predicted richness

## 4. DISCUSSION

Across a 60,000 km^2^ expanse of tropical dry ecosystem in the Eastern Ghats of peninsular India, climate gradients—temperature in particular—played a strong role in plant species distributions and community assembly. For four different life forms, i.e., trees, shrubs, climbers and herbs, a substantial proportion of species showed clear signatures of being associated with either warm-dry or cool-wet conditions. Species with a preference for warm conditions also peaked in at warmer sites, i.e., their niche optima was in a zone of higher than average temperatures for the region. Notably, species with the strongest affiliations to limits of the prevailing temperature conditions had narrower niches than species that peaked in intermediate conditions, suggesting specialization for either cool or warm temperatures among a majority of species. Emerging from species distribution patterns, warm-associated species dominated assemblages at warmer sites and vice-versa. Tree, shrub and climber assemblages in warmer climates had species with wider niches, although tree communities at the warmest limits of the climatic range displayed narrower niches than communities at more intermediate conditions. Herb communities show decreasing niche width at warmer conditions, suggesting specialization in warmer sites.

### 4.1 Patterns in species’ climate niche

Nearly 60% of species, for all life-forms together, showed associations to warm, dry conditions as expected for a tropical dry ecosystem where species would be warm-adapted. Many species showed significantly negative second-order response to climate gradients; for warm-associated species this suggests that they approached their distribution peaks within the conditions found here whereas cool-associated species strongly declined in their probability to occur in warmer conditions. As a result, the most preferred conditions (niche optima) for species strongly reflected their association trend. Warm-associated species had peak occurrence at warmer sites, this pattern being most prominent for trees.

Tree species preferring the cooler parts of the climate gradient had notably narrower niches than species associated to warmer conditions. This agrees with patterns seen in wet forests of the Western Ghats where species associated with drier conditions had wider niches than species in wet, less seasonal conditions (Krishnadas et al., 2021). Whether such spatial segregation of niches emerged from physiological specialization or competitive exclusion arising from trade-offs needs to be determined (Pang et al., 2026). However, species with occurrence optima at either end of the climate gradient had narrow niches, suggesting that being specialized to either end of the climate gradient may compromise a species’ ability to persist in intermediate conditions. In other words, trade-offs may drive tree assembly in the warmest and coolest parts. Thus, some species counter the expectation that species in more seasonal conditions have wider niches on account of having evolved to withstand fluctuating conditions (Quintero & Wiens, 2013).

Similar to trees, shrub and climber species associated with warmer climates had wider niches across the climate space, suggesting that warm-associated species were generalists that persisted into cool, wet sites (Rather et al., 2021). While woody climbers are known to increase in drier, more open conditions (Schnitzer 2005), the pattern for shrubs was contrary to our expectation. It might be that the high temperature and low precipitation levels impose a strong filter and only shrubs that withstand these generally warm, dry conditions can persist here. Our findings reflect observations of shrub species having the capacity to persist across a range of climatic conditions (Naito & Cairns, 2011).

Herbaceous species associated with warmer conditions had their optima there and had narrower niches, indicating specialization for warm-dry conditions. Many herbaceous species, being shade intolerant, may be unable to persist in the coolest, wettest sites with closed-canopy forests, but our sampling scheme of small plots could have missed locally rare species. The trends we found for herbaceous species are likely driven by savanna specialists such as C4 grasses (Nerlekar et al., 2024; Sage & Monson, 1999; Edwards et al., 2010), and corroborate recent findings of high endemism among herbaceous plants in dry ecosystems of India (Nerlekar et al., 2024; Bharathi et al., 2026; Watve, 2013).

Across life-forms, local topography and elevation conveyed no significant additional information as compared to what was already accounted for by climate in explaining species distributions (Supplementary Information, Table S7). For shrubs, herbs, and climbers, climate was the overwhelming driver of species distributions, with climatic models explaining on average between 85% and 90% variance for individual species within each. Nevertheless, in all three life forms, a few species had notable responses to topography, with slope and aspect explaining >50% variance in their distributions. The role of topography could be on account of microhabitat conditions, e.g., valleys tend to be cooler and have higher moisture, ridge tops can be warmer, and steeper slopes can be drier. In TDE, microhabitat may be a key factor for spatial niche partitioning, especially among shallow-rooted shrubs and herbs (Govaert et al., 2024). Analysis from a 4500 km2 subset of this region indeed implicates soil moisture and nutrients as primary drivers of tree community assembly (Rajendra et al., in review).

### 4.2 Species distribution shapes community assembly

Species niche properties propagated to community assembly across the climate gradient. For all life forms, warmer sites unsurprisingly comprised primarily of species associated to and most likely to occur at warmer and drier conditions, suggesting thermal stress as a key filter that organized spatial variation in species composition. Moreover, tree, shrub and climber assemblage at warmer sites all had species with wider niches on average, which indicates nested changes in species composition from cool to warm climates. Such loss of species may accrue from only a subset of species having the physiology tolerance to stressful climates (Spasojevic et al. 2014; Butterfield & Munson, 2016). However, in contrast to shrubs and herbs, tree assemblages had higher species richness at the ends of the climate gradient, which read together with low niche widths at these ends suggest distinct sets of species that have specialized to warm vs cool climates. The comparatively wider niches of tree assemblages at the warmest sites compared to the coolest sites indicates a proportionately greater presence of species that persist across a wider climate niche, lending support to the physiological tolerance hypothesis that should be tested using traits and transplant experiments.

Shrub and herb communities showed highest richness at the intermediate climatic conditions, corroborating filtering at the warm and cool ends. Assemblages with higher predicted richness had species with wider niches, showing that generalist species increased richness in intermediate climate conditions. Shade-adapted shrub and climber species found in the understorey may not withstand higher temperatures or drought in the warm-dry sites (Ehleringer & Mooney, 1983) which could account for the filtration in these communities from cool to warm sites As with species-level patterns, herbaceous species contrasted with other life-forms in assembly across the climate gradient. Even as communities in warmer sites largely comprised warm associated species that preferred warm conditions (niche optima), mean niche width decreased. Assemblages at warmer sites likely consist of dry and warm-adapted C4 grasses (Nerlekar et al., 2024; Sage & Monson, 1999; Edwards et al., 2010), with a turnover of functional groups towards wetter sites where dry-adapted C4 species may be outcompeted by C3 species (Ehleringer & Monson, 1993; Ehleringer et al., 1997). Recent work shows grassland and savanna systems to be underappreciated repositories of herbaceous diversity, in India (Nerlekar et al., 2024) and globally (Nerlekar & Veldman, 2020). Our findings emphasize the need for more detailed surveys of herbaceous plant communities across the Eastern Ghats and a deeper evolutionary understanding of their assembly.

Taken together, the assembly patterns suggest that the physiological tolerance hypothesis largely prevails in TDE for trees, shrubs and climbers, similar to humid forests where it results in nested loss of species shaping community assembly of trees from wet to dry sites (Krishnadas et al., 2021; Esquivel-Muelbert et al., 2017). The contrast from wet ecosystems in the assembly of tree communities at the warmest ends of the TDE may be on account of the relative harshness of climatic conditions (Norden et al., 2026). The driest sites in a wet tropical ecosystem will still be benign compared to the warm sites in a TDE where summer temperatures can reach as high as 45◦C and rainfall can be as low as 500 mm/year. The climatic stress in warm, dry parts of the TDE may allow only those plant species with adequate physiological adaptation to the long seasonal drought and hot summer months, but this specialization may preclude them from tolerating competition in wetter sites. Fire can be an additional selection pressure in drier sites, although we did not measure it explicitly, and hot and dry sites in TDE may be dominated by fire-adapted species.

### 4.3 Conclusion

Temperature and precipitation combinations being the prominent driver of species distributions begs investigation of the role of relevant traits for plant community assembly in tropical dry ecosystems (Sastry et al., 2018). Do warm and cool associated species differ in their hydraulic and/or thermal physiology and do these physiological traits mechanistically link performance and distribution across climate gradients? (Govaert et al., 2021; Kiebacher et al., 2023, Wang et al., 2023). Overall, species’ niche properties across the tropical dry ecosystem of peninsular India implicate hydrothermal conditions as primary controls on plant community assembly, suggesting vegetation sensitivity to shifts in temperature and precipitation. With the caveat that large-scale distributions only capture realized niches, and niche reorganization can depend on biotic interactions, especially at range edges, our results provide a mechanistic basis to project ecosystem responses under changing climate regimes. Linking niche dynamics with landscape scale heterogeneity will enable identification of climate refugia and vulnerability hotspots to plan land-use, restoration, and conservation. Larger-scale censuses and remote-sensing approaches should be combined with smaller-scale monitoring of species performance and community dynamics to guide adaptive, evidence-based policy interventions in tropical dry ecosystems. Multi-scale efforts will be needed to improve ecological forecasting of species populations, community structure and ecosystem function of dry tropical ecosystems in a changing climate.

## Acknowledgements

We thank the National Centre for Biological Sciences (NCBS) and CSIR-Centre for Cellular and Molecular Biology (CCMB) high performance computing clusters for providing computational resources used in this study. We thank Dr. Gaurav Baruah for his comments and recommendations regarding the manuscript and Radhika Rajendra for helping obtain the shapefile used and other inputs and suggestions.

## Data and code availability statement

All data and code used in this article are available publicly on [CaFE Lab / EG Vegetation Patterns · GitLab](https://gitlab.com/thecafelab/eg-vegetation-patterns/-/tree/main). It will be archived using Figshare upon acceptance.

