## Supplementary Material 1 for "Species climate-niche properties suggest that both physiological tolerance and stress dominance shape plant community assembly across a tropical dry ecosystem"

**SUPPLEMENTARY INFORMATION**

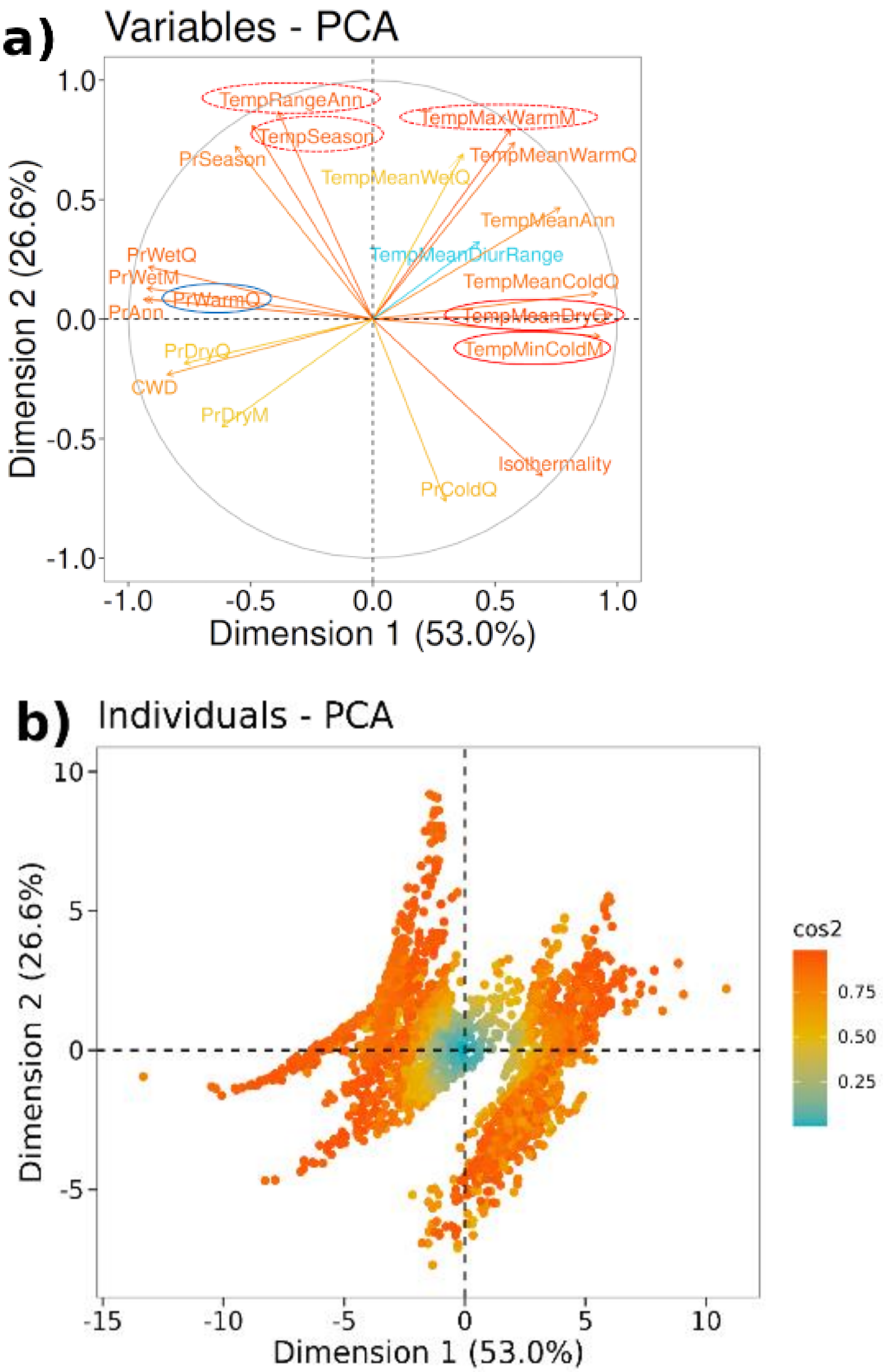

**FIGURE S1** Principal Components Analysis of climate variables. Composite characterization of climatic conditions across the study region using (a) 19 bioclimatic variables from WorldClim 2 (Fick & Hijmans, 2017), to get (b) composite climatic condition for each plot in the PC1 and PC2 space. The colours indicate the strength of representation on the PC axis.
All variables were transformed and scaled before PCA. The variables include Annual Precipitation (PrAnn), Precipitation of Wettest Quarter (PrWetQ), Precipitation of Driest Quarter (PrDryQ), Precipitation of Warmest Quarter (PrWarmQ), Precipitation of Coldest Quarter (PrColdQ), Precipitation of Wettest Month (PrWetM), Precipitation of Driest Month (PrDryM), Precipitation Seasonality (PrSeason), Annual Mean Temperature (TempMeanAnn), Mean Temperature of Warmest Quarter (TempMeanWarmQ), Mean Temperature of Coldest Quarter (TempMeanColdQ), Mean Temperature of Driest Quarter (TempMeanDryQ), Mean Temperature of Wettest Quarter (TempMeanWetQ), Max Temperature of Warmest Month (TempMaxWarmM), Min Temperature of Coldest Month (TempMinColdM), Temperature Annual Range (TempRangeAnn), Mean Diurnal Temperature Range (TempMeanDiurRange), Isothermality, Temperature Seasonality (TempSeason).

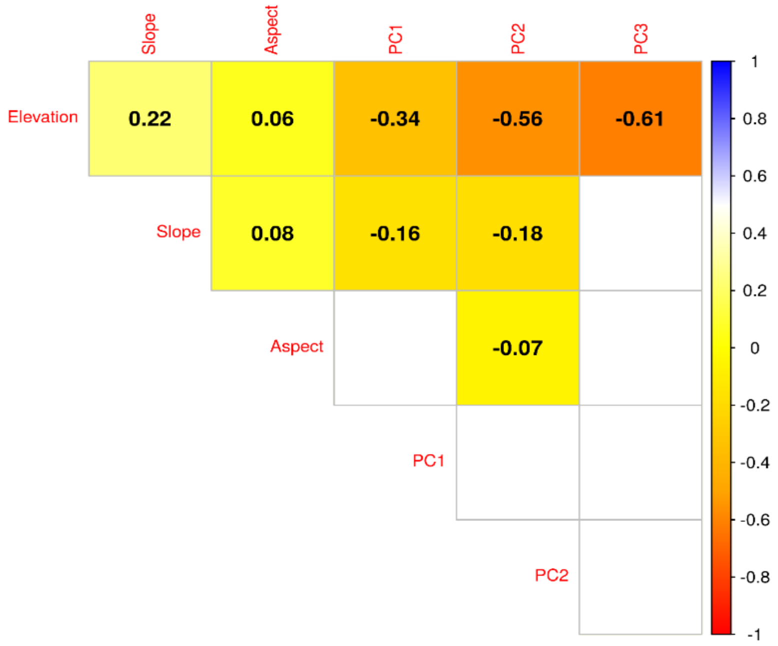


**FIGURE S2**. Correlation between climate and topography. For 16 variables downloaded from WorldClim and three topological variables (elevation, slope, and Aspect. The numbers represent the Pearson's correlation coefficient with the colours representing a similar information visually. The blank cells have correlations that are not statistically significant at $P < 0.05$

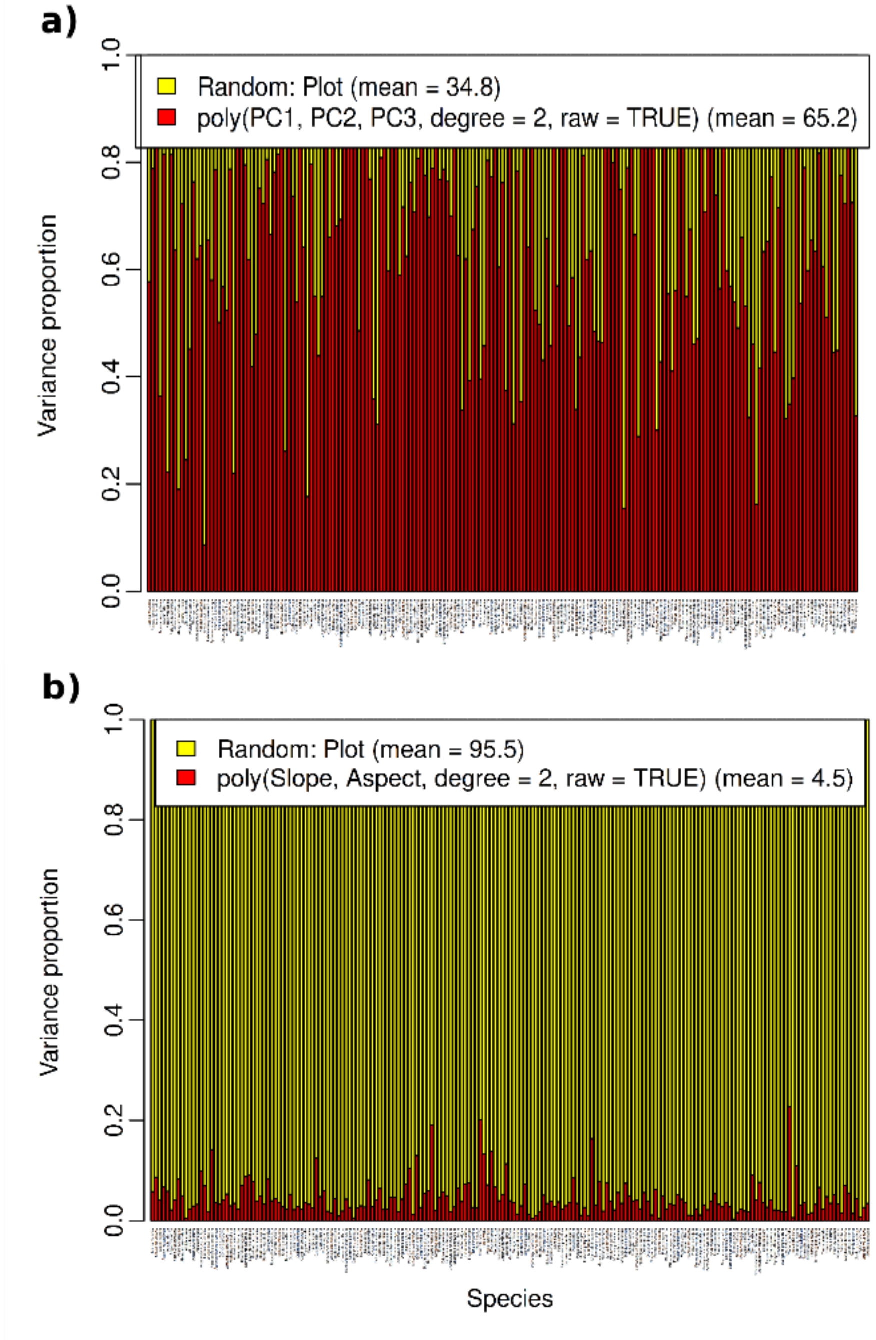


**FIGURE S3.** Proportions of variance explained by models using (a) Climate and (b) Topography for each tree species on the x-axis. The polynomial functions representing the total effects of all the climatic PCs explain on average 65 percent of variance in tree species distribution, in contrast to only 4.5 percent by the topographical variables. Random indicates baseline variation in occurrence not attributable to the variables included in the model.

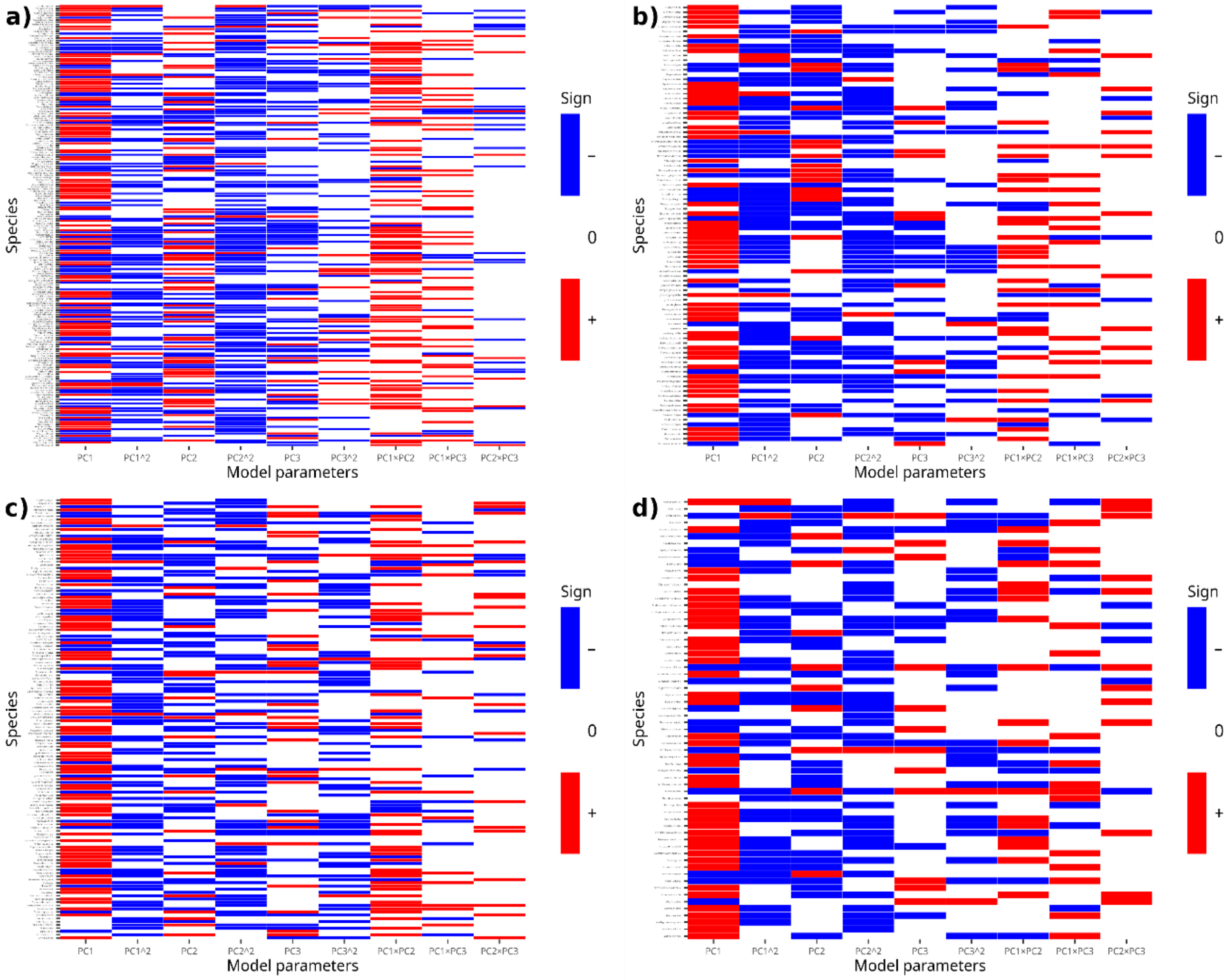


**FIGURE S4** Species association with the different levels of the climatic model and their interactions for (a) trees, (b) shrub, (c) herb, and (d) climber. The first and second order of PC1, PC2, and PC3 as well as interactions of their first order components are shown. Red and blue colour indicate positive and negative associations, respectively, of species to the corresponding predictor. The patterns reveal a predominantly positive association of species to PC1 axis across all life forms, i.e., an increase in occurrence at warm-dry sites across the region, which was to be expected for species in a tropical dry ecosystem.

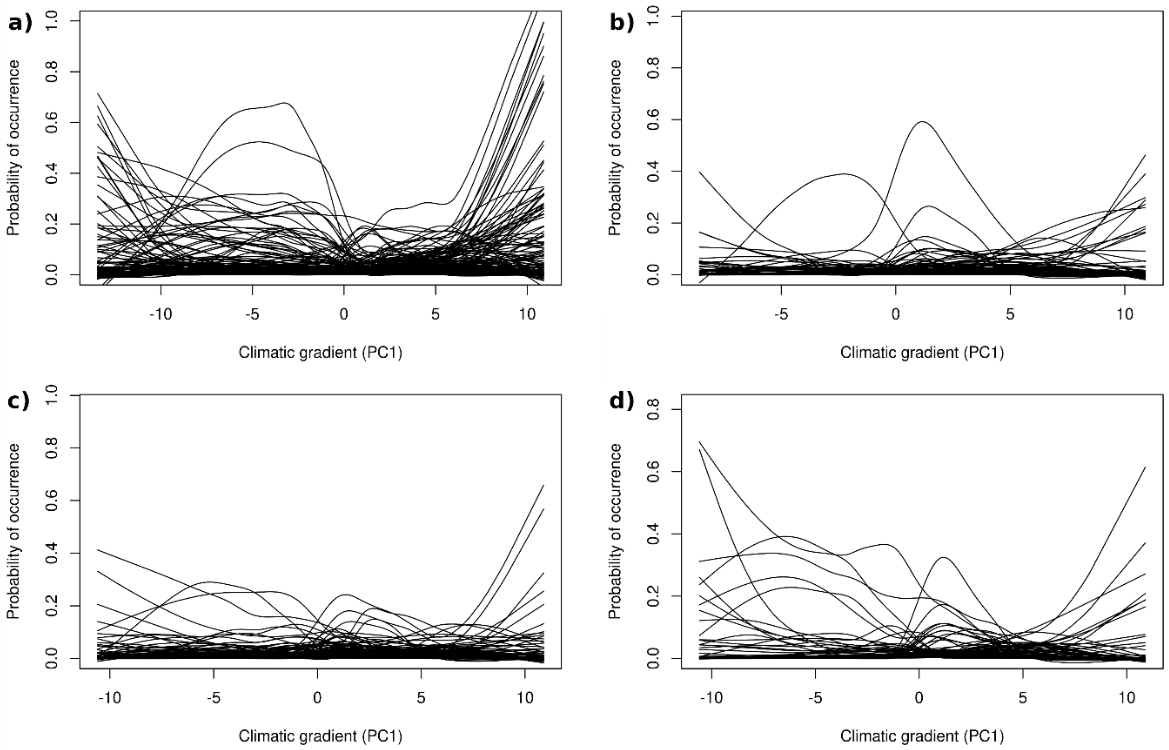


**FIGURE S5.** Predicted probability of occurrence for each species across the climatic gradient for (a) tree, (b) shrub, (c) herb, and (d) climber. Shrubs, herbs and climbers had relatively higher probability of occurrence than trees in the intermediate range of climate than the climatic extremes.

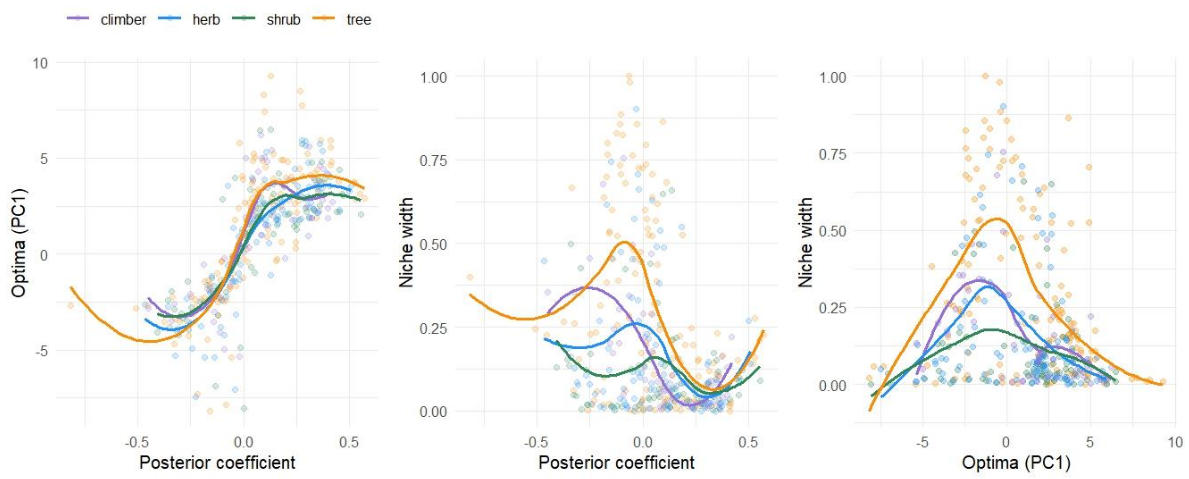


**FIGURE S6.** Species' level relationships between niche properties when niches are estimated based on median occurrence probability at a site, without considering uncertainty (i.e., not based on lower limit of confidence interval). Compare to analyses where species are uncluded only when the lower limit of the confidence interval of estimates does not overlap zero, here 367 out of 485 species get included. Nevertheless, trends remain strongly similar (see main Fig. 2).

| Model | No. of species | Mean AUC | Median AUC | Mean Tjur's $R^2$ | Median Tjur's $R^2$ | Mean RMSE |
|---|---|---|---|---|---|---|
| Climber Clim | 64 | 0.870 | 0.891 | 0.076 | 0.039 | 0.150 |
| Climber Topo | 64 | 0.853 | 0.867 | 0.058 | 0.023 | 0.153 |
| Herb Clim | 136 | 0.898 | 0.913 | 0.082 | 0.052 | 0.132 |
| Herb Topo | 136 | 0.891 | 0.917 | 0.074 | 0.034 | 0.131 |
| Shrub Clim | 92 | 0.887 | 0.891 | 0.073 | 0.045 | 0.136 |
| Shrub Topo | 92 | 0.892 | 0.910 | 0.066 | 0.032 | 0.135 |
| Tree Clim | 193 | 0.904 | 0.905 | 0.117 | 0.090 | 0.159 |
| Tree Topo | 193 | 0.915 | 0.918 | 0.117 | 0.089 | 0.158 |

**TABLE S7.** Model fit statistics of HMSC models employed-Area under curve, Tjur's $R^2$ and RMS error. For each life form (climber, herb, shrub, tree) two models (climatic and topographic) were run. Fit for the two types of models is high and very similar in each case, implying no additional information is obtained from topographic models and climate alone explains a large proportion of variability in the data.
